# Full-length immunoglobulin high-throughput sequencing reveals specific novel mutational patterns in POEMS syndrome

**DOI:** 10.1101/824722

**Authors:** Sébastien Bender, Vincent Javaugue, Alexis Saintamand, Maria Victoria Ayala, Mehdi Alizadeh, Matthieu Filloux, Virginie Pascal, Nathalie Gachard, David Lavergne, Fabienne Auroy, Michel Cogne, Frank Bridoux, Christophe Sirac, Arnaud Jaccard

## Abstract

POEMS syndrome is a rare multisystem disease due to an underlying plasma cell (PC) dyscrasia. The pathophysiology of the disease remains unclear but the role of the monoclonal immunoglobulin (Ig) light chain (LC) is strongly suspected, due to the highly restrictive usage of two λ variable (V) domains (IGLV1-40 and IGLV1-44) and the general improvement of clinical manifestations following PC clone-targeted treatment. However, the diagnostic value of Ig LC sequencing, especially in case of incomplete forms of the disease, remains to be determined. Using a sensitive high-throughput Ig repertoire sequencing on RNA (RACE-RepSeq), we detected a λ LC monoclonal expansion in the bone marrow (BM) of 85% of patients with POEMS syndrome, including some in whom bone marrow tests routinely performed to diagnose plasma cell dyscrasia failed to detect λ^+^ monoclonal PCs. Twenty-four of the 30 LC clonal sequences found (80%) were derived from the IGLV1-40 and IGLV1-44 germline genes, two from the closely related IGLV1-36 gene, and all were associated with an IGLJ3*02 junction (J) gene, confirming the high restriction of VJ region usage in POEMS syndrome. RACE-RepSeq VJ full-length sequencing additionally revealed original mutational patterns, the strong specificity of which might crucially help establish or eliminate the diagnosis of POEMS syndrome in uncertain cases. Thus, RACE-RepSeq appears as a sensitive, rapid and specific tool to detect low-abundance PC clones in BM, and assign them to POEMS syndrome, with all the consequences for therapeutic options hereby.

## Introduction

Polyneuropathy, organomegaly, endocrinopathy, monoclonal gammopathy, and skin changes (POEMS) syndrome is a multisystem disorder. Apart from the above manifestations, sclerotic bone lesions, edema, anasarca, thrombocytosis, and various other symptoms are common. Diagnosis relies on the presence of at least three major criteria and one minor criteria.^1^ Amongst major criteria, monoclonal plasma cell (PC) disorder and polyradiculoneuropathy are mandatory. Other major criteria include bone lesions, high level of vascular endothelial growth factor (VEGF) and Castleman disease. Minor criteria are numerous, reflecting the heterogeneity of the disease manifestations; they include organomegaly, endocrinopathy, skin changes, papilledema, extravascular volume overload, and thrombocytosis.

POEMS syndrome has been recently incorporated within the spectrum of the monoclonal gammopathies of clinical significance.^2^ Indeed, even if the pathophysiology of the disease is yet unknown, an efficient treatment of the underlying PC clone usually results in improvement of all manifestations. The clones are often small^3,4^ and almost always characterized by the production of a λ light chain (LC) deriving from any of two variable (V) germline genes, IGLV1-40 or IGLV1-44.^5,6^ Consequently, it has been suggested that, through still unknown pathways, the monoclonal LC structure might specifically trigger the major biologic anomaly suspected to account for the POEMS pathophysiology: VEGF hyperproduction. High VEGF levels indeed appear to account for most features of the disease.^7,2^ Immunochemically, a small free λ LC component is virtually always associated with a complete immunoglobulin, most frequently IgAλ, or IgGλ. Rarely, the λ LC is the sole monoclonal component. Hypergammaglobulinemia is frequent in POEMS syndrome and due to the increase in polyclonal κ free LCs, nearly 80 % of the cases presents with a normal κ/λ ratio despite slight increase in serum monoclonal λ free LC level.^8,9^ Treatment is conditioned by the characteristics of the underlying PC clone. Radiotherapy is used for localized plasmacytoma, whereas systemic chemotherapy is required in patients with bone marrow (BM) infiltration or with too many or no bone lesions.^1,10^ Thus, detection and identification of a BM PC clone is a crucial step of management. To date, a PC clone is found in up to 66% of patients with POEMS syndrome using flow cytometry, immunohistochemistry, and/or in situ hybridization technics.^11^ However, this percentage may be either underestimated due to low level PC infiltration, especially in the presence of a concomitant plasmacytoma,^12^ or overestimated when non-relevant clones are present in the bone marrow. In this setting, due to the homogeneity of Ig V domain usage,^5,6^ Ig sequencing may be of great value to determine the pathogenicity of a detected clone, particularly in patients with atypical clinical presentation.

Next generation sequencing (NGS) has changed the paradigm of Ig repertoire analysis and detection of minimal residual disease (MRD) in B cell malignancies, including multiple myeloma (MM).^13^ Kawajiri-Manako C *et al*. showed the possibility to use NGS to identify PC clone LC rearrangements on DNA extracted from the BM of patients with POEMS syndrome.^12^ However, due to the low clonal expansion of PCs in BM, they detected clonal λ LC rearrangements in only 11 out of 30 patients. Moreover, the PCR amplification design usually used for characterizing the Ig repertoire on DNA only encompasses the CDR3 region. While this strategy might give a good evaluation of IgH VDJ genes, it can be expected with a poor efficacy when evaluating the clonality of light chain genes (which carry little of absent N insertions at the V-J region, and which usually display more canonical rearrangements than VHDJH genes with poorly diversified CDR3 length). This strategy of characterizing clones through a short CDR3 stretch instead of the full VJ regions additionally overlooks most of the clonal diversification related to somatic hypermutation (thus occulting intra-clonal diversity), and might even be unable to assign a VJ rearrangement to one or the other member of a VL subgroup (notably in the case of the closely related V germline genes IGLV1-44 and IGLV1-36).^12^ By contrast, we propose herein a method for high-resolution characterization of the full-length VJ region. This relies on a sensitive and rapid cDNA repertoire analysis of BM or bone lesions/plasmacytoma through high-throughput sequencing, in order to secure diagnostic approach and management in POEMS syndrome.^14^

## Methods

### RNA sample preparation

Total RNAs were extracted and prepared from bone marrow aspirates or biopsies or plastocytoma biopsies using TRIzol reagent (Ambion).

### TOPO cloning and sequencing

Complementary DNAs were obtained by reverse transcription performed using the High Capacity cDNA Archive Kit (Applied Biosystems) and 1μg of total RNAs. Polymerase chain reaction (PCR) amplification was performed using a 3’ consensus primer complementary to all λ constant regions (5’ CTCCCGGGTAGAAGTCACT 3’) and a 5’ consensus primer for all Vλ subgroups leader regions (5’ ATGGCCTGGDYYVYDCTVYTYCT 3’). PCR products were then cloned into pCR2.1 TOPO vector (Invitrogen), and DNA sequencing was carried out using Big-Dye terminators (Applied Biosystems) on a 16-capillaries electrophoresis system 3130 XL (Applied Biosystems). Sequences were analyzed using FinchTV software and aligned with MultAlin (http://multalin.toulouse.inra.fr/multalin/) and the λ VJ rearrangement was determinated using IMGT/V-QUEST tool (http://imgt.org/).^15–17^

### High-Throughput sequencing on ROCHE 454 sequencer

Transcripts were amplified using 5’ RACE-PCR and consensus reverse primers hybridizing in λ constant exons as previously described^18,19^ and sequenced on a 454 sequencer (Roche). Repertoire analysis was done using IMGT/HIGHV-QUEST tool (http://imgt.org/).^18^

### High-Throughput sequencing on Illumina MiSeq sequencer

For each sample, up to 2μg of total RNAs were used for high-throughput sequencing.

#### 5’Rapid Amplification of cDNA Ends (RACE) PCR

First, a mix containing up to 2μg of RNAs, 1μL of a consensus reverse primer of all lambda constant regions (5’CTGGCCGCYTACTTGTTGTT 3’) and 1 μL of a dNTP solution mix (New England Biolabs), adjusted to a final volume of 12 μL with water, was incubated 3 min at 72°C and 2 min at 42°C. After a short spin, the mix was placed on ice for 2 min. Then, 4 μL of ProtoScript buffer, 2μL of DTT solution, 1μL of ProtoScriptII (New England Biolabs) and finally 1 μL of a cap-race primer (5’ AAGCAGTGGTATCAACGCAGAGTACAT(GGGG) 3’, the 4 G between brackets are ribonucleotides) were added. This mix was incubated 90 min at 42°C, 10 min at 70°C and then stored at 4°C. At the end of the reaction, 20μL of RNAse free water were added.

#### Library preparation

The λ cDNA were amplified with Taq Phusion (New England Biolabs) using an universal forward universal primer mix (5’ CTAATACGACTCACTATAGGGC 3’ and 5’ CTAATACGACTCACTATAGGGCAAGCAGTGGTATCAACGCAGAGT 3’, in a ratio of 4:1) as described^20^ and a reverse primer hybridizing within the λ constant exons (5’ GTCTCGTGGGCTCGGAGATGTGTATAAGAGACAGGTTGGCTTGYAGCTCCTCAG 3’). Four PCR replicates of 25 μL were done. Cycling conditions were 30 sec at 98°C, 32 cycles of 30 sec at 98°C, 30 sec at 65°C, 30 sec at 72°C and final elongation 5 min at 72°C. The four replicates were pooled and migrated in a 2% agarose gel (alternatively, Ampure XP beads can be used). The band of interest (±550 pb) was cut and the DNA was eluted in 30 μl of elution buffer using the kit Nucleospin gel and PCR clean-up (Macherey-Nagel). Then, Illumina sequencing adapter and tag sequences were added by primer extension. For this, previous DNA were reamplified with Taq Phusion. Four PCR replicates of 25 μL were made using primers containing a unique couple of tag per sample (a tag is a unique combination of 8 nucleotides). Cycling conditions were 30 sec at 98°C, only 12 cycles of 30 sec at 98°C, 30 sec at 65°C, 30 sec at 72°C and final elongation 5 min at 72°C. The four replicates were mixed and another migration in a 2% agarose gel was done (alternatively, Ampure XP beads can also be used). The band of interest (±600 pb) was cut and DNAs were eluted twice in 30 μL of elution buffer using the kit Nucleospin gel and PCR clean-up (macherey-Nagel).

#### Sequencing on Illumina MiSeq and repertoire analysis

Resulting amplicons were sequenced on an Illumina MiSeq sequencing system using MiSeq Reagent kit V3 600 cycles (alternatively, MiSeq reagent kit V2 can be used but with lower cluster density and shorter sequences). Paired reads were merged using FLASH.^21^ Repertoire analysis was done using the VIDJIL tool (http://vidjil.org/)^22^ (Supplemental Figure 1) and IMGT/HIGHV-QUEST tool (http://imgt.org/).^18,20^

### Sequencing of IGLV genes on DNA

DNA extraction from a lymph node biopsy and sequencing of IGLV gene were done using the EuroClonality/BIOMED-2 guidelines as previously described^23,24^ with an IGLV forward consensus primer (5’ ATTCTCTGGCTCCAAGTCTGGC 3’) specific to the FR3 region and an IGLJ reverse consensus primer (5’ CTAGGACGGTGAGCTTGGTCCC 3’).

## Results

A total of 36 samples from patients with confirmed POEMS syndrome, were analyzed by Ig RNA sequencing, mainly with high-throughput sequencing (RACE-RepSeq) except for four patients that were characterized only with TOPO cloning and Sanger sequencing (p1, p3, p6, p21). Twenty-four patients (67%) had bone lesions and fifteen patients (42%) had BM monoclonal plasma cell infiltration identified by immunohistology on BM biopsy aspiration (Table 1). Bone marrow samples were available in 33 patients. The remaining samples were obtained from bone lesions in 3 cases, (p7, p8, p18). In one patient, both biopsy samples from BM and from a bone lesion were analyzed (p15). In another patient with an atypical form of POEMS syndrome (p33), both biopsy samples from BM and from a lymph node were analyzed. Ig RNA sequencing allowed finding an IGLV clone in all patients with a detectable BM PC clone (Table 1). Interestingly, a clonal IGLV gene was found in 13 out of 19 BM samples with negative cytology and cytometry or with monotypic κ PCs (Table 1). All but one of these patients presented with one or several plasmacytomas. A comparison of the TOPO cloning method with our RACE-RepSeq method revealed that 7/11 samples previously negative for IGLV clone using Sanger sequencing were subsequently found positive by NGS. Overall, we detected at least one IGLV clone in 31/36 patients (86%) and in 27/33 patients (82%) whose BM samples were analyzed. For patient 15, the same monoclonal LC sequences were detected in both BM and bone lesion samples (Supplemental Figure 1). Concerning patient 33, with an atypical form of POEMS syndrome (polyneuropathy, circulating monoclonal IgGλ, elevated serum VEGF but no bone lesion or monoclonal plasmocytosis in the BM and multiple lymph nodes with an aspect of follicular hyperplasia with monoclonal λ^+^ PC), the BM was negative by RACE-RepSeq but we detected an IGLV monoclonal sequence in a lymph node biopsy (included in paraffin) obtained from a Sanger DNA sequencing surrounding the V-J junction (CDR3) only.

**Table 1 :** Clinical, biological and monoclonal λ LC characteristics of POEMS syndrome patients.

| Patient Id | Bone marrow | Plasmacytoma | SANGER analysis | NGS analysis | IGLV gene | IGLJ gene | Mutation | Monoclonal Ig | [λFLC] (at diagnosis) | [VEGF] (at diagnosis) | IGHV gene |
| --- | --- | --- | --- | --- | --- | --- | --- | --- | --- | --- | --- |
| p4 | Negative | yes | Clone found | Clone found (Roche) | IGLV1-44 | IGLJ3*02 | 38→P | $\alpha\lambda$ | 15.8 | 1618 | IGHV3-11 and IGHV4-39 |
| p6 | Negative | yes | Clone found | ND | IGLV1-44 | IGLJ3*02 | 38→P | $\gamma\lambda$ | 10.8 | 420 | |
| p9 | Negative | yes | No clone | Clone found | IGLV1-44 | IGLJ3*02 | 38→P | $\gamma\lambda$ | 23.3 | 162 | other clone (IGLV3-1/IGLJ2*01) |
| p10 | Negative | yes | ND | Clone found | IGLV1-44 | IGLJ3*02 | 38→P | $\alpha\lambda$ | 5.78 | 430 | IGHV3-9 |
| p11 | Negative | yes | ND | Clone found | IGLV1-44 | IGLJ3*02 | 38→P | $\gamma\lambda$ | 100 | 3752 | |
| p13 | Negative | yes | ND | Clone found | IGLV1-44 | IGLJ3*02 | 38→P | $\alpha\lambda$ | 161 | 63332 | |
| p14 | Negative | yes | ND | Clone found | IGLV1-44 | IGLJ3*02 | 38→P | $\alpha\lambda$ | 32 | 2300 | |
| p17 | Negative | no | ND | Clone found | IGLV1-44 | IGLJ3*02 | 38→N | $\alpha\lambda$ | - | 2000 | |
| p1 | Positive | yes | Clone found | ND | IGLV1-44 | IGLJ3*02 | 38→P | $\alpha\lambda$ | 26.3 | 1965 | IGHV4-38-2 |
| p2 | Positive | yes | Clone found | Clone found (Roche) | IGLV1-44 | IGLJ3*02 | 38→P | $\gamma\lambda$ | 104 | 1082 | IGHV2-26 |
| p3 | Positive | yes | Clone found | ND | IGLV1-44 | IGLJ3*02 | 38→P | $\alpha\lambda$ | 35.4 | 7000 | |
| p5 | Positive | no | No clone | Clone found (Roche) | IGLV1-44 | IGLJ3*02 | 38→P | $\alpha\lambda$ | 217 | 2725 | |
| p12 | Positive | no | No clone | Clone found | IGLV1-44 | IGLJ3*02 | 38→S | $\alpha\lambda$ | 45 | 7800 | |
| p15 | Positive | yes | ND | Clone found* | IGLV1-44 | IGLJ3*02 | 38→A | $\alpha\lambda$ | 72 | 9055 | |
| p16 | Positive | no | ND | Clone found | IGLV1-44 | IGLJ3*02 | 38→A | $\gamma\lambda$ | 51.9 | 739 | |
| p23 | Negative | yes | No clone | Clone found | IGLV1-40 | IGLJ3*02 | 40→N | No monoclonal Ig | 46 | 845 |  |
| p24 | Negative | yes | ND | Clone found | IGLV1-40 | IGLJ3*02 | 40→N | - | 47 | 1300 |  |
| p20 | Negative | yes | No clone | Clone found | IGLV1-40 | IGLJ3*02 | 40→N | $\gamma\lambda$ and $\gamma\kappa$ | 150 | 1400 | |
| p19 | Positive | no | ND | Clone found | IGLV1-40 | IGLJ3*02 | 40→N | $\alpha\lambda$ | 43.3 | 11188 | |
| p21 | Positive | yes | Clone found | ND | IGLV1-40 | IGLJ3*02 or 01 or IGLJ2*01 | 40→Q | $\gamma\lambda$ | 134 | 1423 | |
| p22 | Positive | no | ND | Clone found | IGLV1-40 | IGLJ3*02 | 40→N | $\alpha\lambda$ | 32 | 7290 | IGHV2-70 |
| p25 | Positive | no | ND | Clone found | IGLV1-36 | IGLJ3*02 | - | $\gamma\lambda$ | 37.8 | 5000 | other clone (IGLV2-14/IGLJ3*02) |
| p26 | Positive | no | ND | Clone found | IGLV1-36 | IGLJ3*02 | - | No monoclonal Ig | 107 | 7550 |  |
| p29 | Negative | yes | ND | Clone found | IGLV3-1 | IGLJ2*01 | - | $\gamma\lambda$ | 40.2 | 3275 | |
| p30 | Negative | no | No clone | Clone found | IGLV3-25 | IGLJ1*01 | - | $\gamma\lambda$ | - | 1966 | |
| p27 | Positive | yes | No clone | Clone found | IGLV3-1 | IGLJ3*01 | - | $\gamma\lambda$ | 35 | 1210 | |
| p28 | Positive | no | Clone found | Clone found | IGLV3-1 | IGLJ2*01 | - | $\alpha\lambda$ and $\gamma\lambda$ | ND | 705 | |
| p31 | Negative | yes | No clone | No clone | No clone | No clone | - | No monoclonal Ig | 41 | 1700 |  |
| p32 | Negative | yes | No clone | No clone | No clone | No clone | - | $\gamma\lambda$ | 51.3 | 1190 | |
| p33 | Negative | no | No clone | No clone | No clone | No clone | - | $\gamma\lambda$ | 25.2 | 1146 | |
| p34 | Negative | yes | No clone | No clone | No clone | No clone | - | $\gamma\lambda$ | 21.8 | 705 | |
| p35 | Negative | yes | ND | No clone | No clone | No clone | - | $\alpha\lambda$ | - | 2550 | |
| p36 | Negative | no | No clone | No clone | No clone | No clone | - | $\alpha\lambda$ | 27 | 665 | |

\* Sequencing on both bone marrow and plasmacytoma, ND=Not Done

|  |  |  |  |  |  |  |  |  |  |  |  |
| --- | --- | --- | --- | --- | --- | --- | --- | --- | --- | --- | --- |
| p7 | Negative | yes | Clone found | Clone found | IGLV1-44 | IGLJ3*02 | 38→A | $\gamma\lambda$ | 15,3 | 1419 | |
| p8 | Negative | yes | Clone found | Clone found | IGLV1-44 | IGLJ3*02 | 38→A | $\gamma\lambda$ and $\alpha\kappa$ | 55,6 | 3560 | |
| p18 | Positive | yes | ND | Clone found | IGLV1-44 | IGLJ3*02 | 38→P | $\gamma\kappa$ and $\gamma\lambda$ | 21,6 | 1100 | |

Bone marrow sequencingPlasmacytoma sequencing

In accordance with previous publications,^5,6^ IGLV1-40 or IGLV1-44 variable genes were overrepresented and identified in 25 of all patients (69 %), and in 81% of patients in whom a λ-positive clone was detected by sequencing (25/31). IGLV1-40 or IGLV1-44 monoclonal sequence was found in the 4 bone lesion biopsy samples. In six patients (17%), other IGLV genes were found, including two patients with a monoclonal IGLV1-36 variable gene, which is closely related to the IGLV1-44 germline gene (Table 1). Finally, no monoclonal λ LC was found in the BM of six patients (17%). Among them, none had a detectable PC clone using conventional cytologic and immunophenotypic analysis, but we finally detected in patient 33 (atypical form of POEMS) an IGLV1-44/IGLJ3*02 clonal sequence in a paraffin embedded lymph node biopsy using Sanger sequencing on DNA (No RNA available from this sample). Ig RNA sequencing allowed the detection of an IGLV1-40 or IGLV1-44 monoclonal LC in the BM from 11 patients (58 %) whose BM samples were previously negative with usual cytological analysis. Among these, 10 were analyzed by NGS (RACE-RepSeq), including three patients previously found negative with Sanger analysis. For two other patients with negative routine/conventional analysis, another λ LC clone was found. All the IGLV1-40, IGLV1-44 and IGLV1-36 variable genes were rearranged on the same IGLJ3*02 joining segment, as previously described.^6^ However, this rearrangement does not lead to a canonical CDR3, since CDR3 sequences were highly diverse (Figure 1). Interestingly, all the other clonal IGLV found, not related to the IGLV1-40, IGLV1-44 or IGLV1-36, were not associated to the IGLJ3*02 gene (Table 1). Finally, we sequenced IGHV genes in five patients and we did not find any homogeneity or recurrent mutations (all five deriving from different IGHV germline genes) (Table 1).

**Figure 1 :**
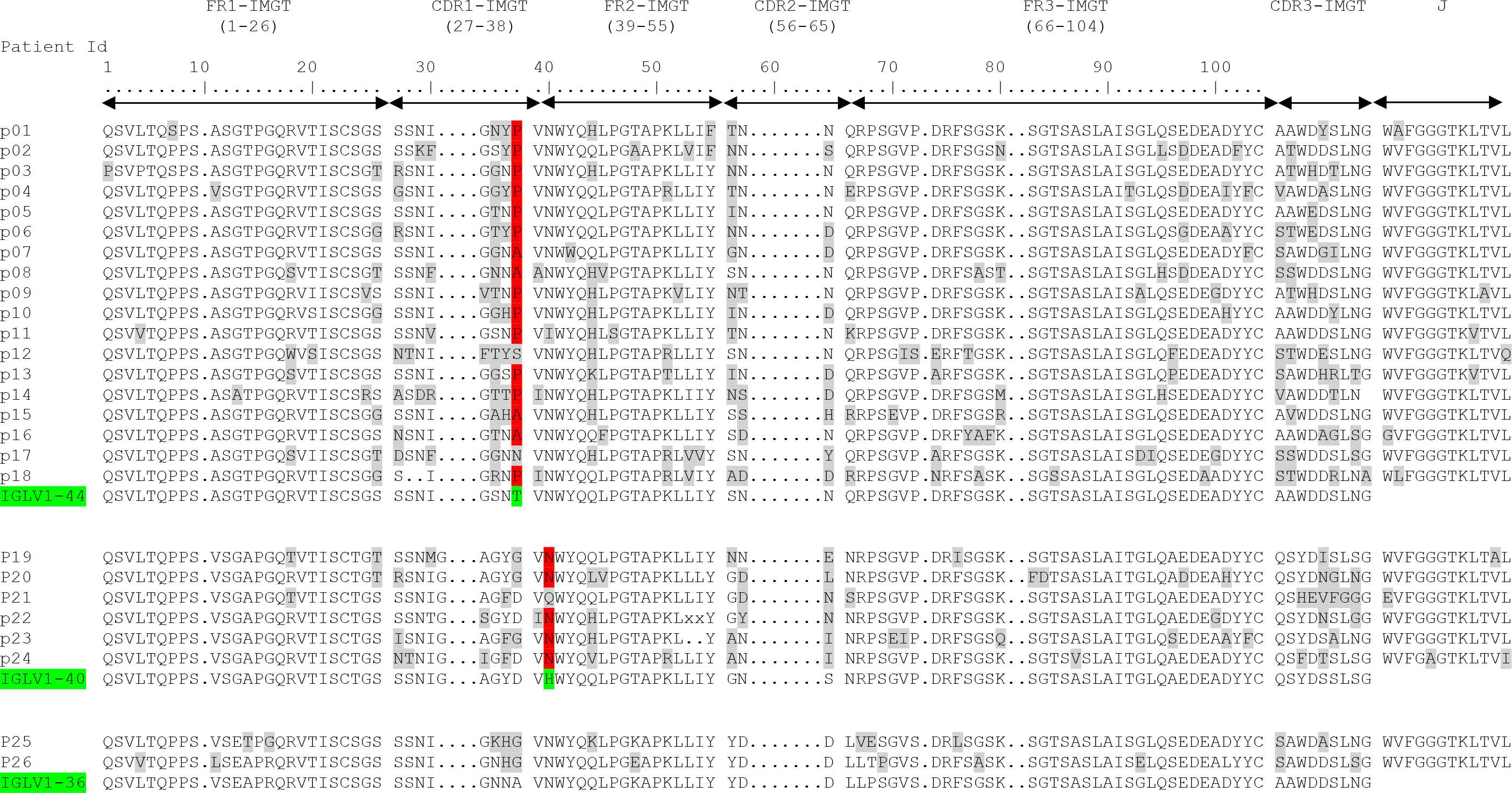
Deduced amino acid sequences of the monoclonal λ LC VJ domains in POEMS syndrome patients compared to germline sequences according to IMGT numbering. Mutated amino acids are highlighted in grey and the redundant mutations in position 38 for IGLV1-44 and position 40 for IGLV1-40 are highlighted in red.

When compared to the corresponding germline sequences, the analysis of the IGLV1-40 or
IGLV1-44 amino-acid (AA) sequences revealed a novel specific mutational pattern with two redundant mutations at position 38 for IGLV1-44 and 40 for IGLV1-40 AA sequences, according to IMGT unique numbering^25^ (Figure 1). 18/18 (100%) of IGLV1-44 sequences are mutated in position 38 and the polar Threonine 38 was replaced by a hydrophobic residue in 16/18 sequences. A proline, which beside its hydrophobicity is known to disrupt protein secondary structures, was found in 12 of the 18 IGLV1-44 sequences (67%) and an alanine in four other cases (22%) leading to a (P/A)VNWYQ consensus stretch in 61% of cases. Interestingly, a retrospective analysis of the series by Abe *et al.* confirms the recurrence of this mutation, with 9/9 patients having a mutation in position 38 including four patients with a proline and one patient with an alanine^6^ (Figure 2). Since IGLV1-44 is frequently associated with AL amyloidosis, we analyzed sequences from previous studies by Perfetti et al.,^26^ Comenzo et al.,^27^ the Boston university AL-Base,^28^ and our own unpublished sequences leading to a total of 100 IGLV1-44 sequences from patients with AL amyloidosis and 6 from other PC disorders. Strikingly, only 4 sequences presented a T→P/A mutation in position 38 (0.04%) and only one reconstituted the full (P/A)VNWYQ stretch (supplemental figure 2). However, this V domain was not associated to the recurrent IGLJ3*02 domain. Regarding our IGLV1-40 AA sequences, the histidine 40 was replaced by an asparagine in five out of six cases (83%), and in the remaining case, the histidine was replaced by a glutamine. Interestingly, the asparagine residue in position 40 is present in the germline gene of the IGLV1-44 and IGLV1-36 domains and reveals a consensus VNWYQ stretch found in the FR2 of 73% of the IGLV1-36, IGLV1-40 and IGLV1-44 sequences (Figure 1). The H40→N substitution in the FR2 region in patients with IGLV1-40 clone is associated with a glutamine or aspartate in position 38 to form a (D/G)VNWYQ stretch that we also found in the 2 patients with IGLV1-36 clonal sequences. This mutation was retrospectively found in all monoclonal IGLV1-40 sequences (n=4) from the series of Martin *et al*. and Abe *et al*. (Figure 2)^5,6^ but was observed in only 1 of the 18 IGLV1-40 sequences (0.06%) from AL amyloidosis and other PC disorders extracted from the AL-Base^28^ and our own database and none reconstituted the full (D/G)VNWYQ consensus stretch.

**Figure 2 :**
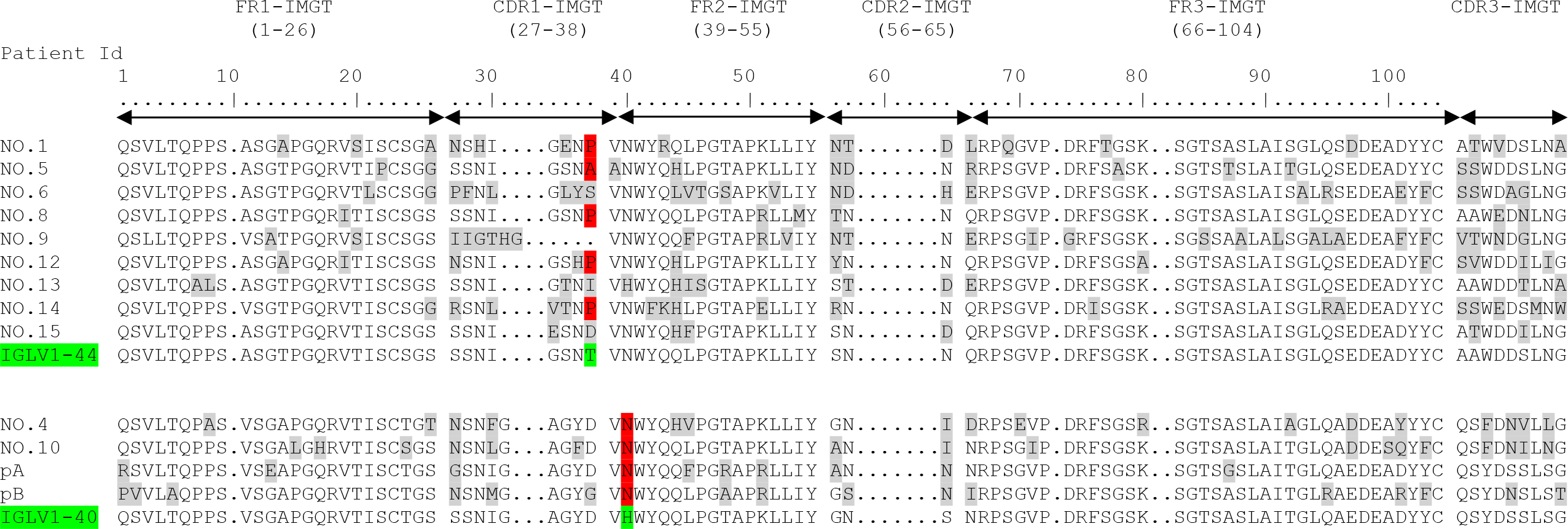
Previously published amino acid sequences of POEMS syndrome compared to germline sequences according to IMGT numbering (NO.1 to NO.15 from Abe et al^6^ and pA/pB from Soubrier et al^5^). Mutated amino acids are highlighted in grey and the redundant P or A mutations in position 38 for IGLV1-44 and N in position 40 for IGLV1-40 are highlighted in red.

## Discussion

Despite the establishment of international criteria,^1,3^ the diagnosis of POEMS syndrome can be challenging, due to the variety of clinical symptoms and frequent atypical clinical presentation lacking the full-blown spectrum of manifestations. Among these, even the main manifestation of the disease, polyneuropathy, is sometimes absent during early disease stage, and the level of vascular endothelial level (VEGF), one of the main diagnostic criteria, may remain within normal range because of previous treatment by intravenous immunoglobulins or corticosteroids. In these situations, the diagnosis of POEMS is often delayed, resulting in delayed treatment introduction and increased morbidity. Indeed, the treatment of POEMS syndrome is highly specific and should be started before the apparition of a severe polyneuropathy. In the present study, we show that LC gene sequencing, particularly using the novel RNA-based NGS technique, is a potent tool to secure the diagnosis of POEMS including patients with atypical presentation, and to guide therapeutic management.

In patients with plasmacytoma but negative BM using conventional analysis or in those lacking mandatory criteria for the diagnosis of POEMS, the search for a PC clone in the BM is crucial to deliver appropriate therapy. Based on the present series of POEMS sequences, we confirm the restriction of λ V domain usage to IGLV1-40 or IGLV1-44 rearranged to IGLJ3*02 gene, as already shown by others.^5,6^ We also found two patients with IGLV1-36 variable gene, and this gene could potentially be a new IGLV1 gene associated with POEMS syndrome, since it is known as a paralog of IGLVλ1-44 gene^29,12^ and contains the consensus sequence VNWYQ in the FR2 domain. Accordingly, an IGLV1-36 sequence associated with POEMS syndrome was recently found by RNAseq in the series of Nagao *et al.*^30^ Another hypothesis could be that the LC hypermutations that appear during B cell response could be responsible for the assignment to an IGLV1-36 derived gene in place of the IGLV1-44 one. In any case, the association of this new λ V domain with POEMS syndrome remains to be confirmed with more sequencing data from POEMS patients. Four other patients had unrelated IGLV clones (IGLV3-1 and IGLV3-25, none of them rearranged with the canonical IGLJ3*02 segment), that probably corresponded to incidental plasma cell clones not responsible for POEMS syndrome. Patients with POEMS syndrome usually display hypergammaglobulinemia, increased polyclonal plasma cells in the bone marrow and frequently more than one monoclonal Ig.^11^ This was confirmed in our series, in which some patients had two concomitant IGLV clones and three were diagnosed with a κ LC clone in the BM (Table 1).

In 2018, Kawajiri-Manako C *et al.* showed for the first time that NGS may be useful to detect IGLV clones in BM from patients with POEMS.^12^ However, NGS-based Ig sequencing on DNA suffers from the frequent low PC burden observed in POEMS syndrome and more generally, in monoclonal gammopathies of clinical significance (MGCS).^2^ Accordingly, significant clonal rearrangements were found in only 11/30 patients and detection of some lambda PC clones required heteroduplex analysis on cDNA.^12^ Our method proposes a sensitive assay based on cDNA analysis of Ig repertoire by NGS (RACE-RepSeq) allowing in addition, the sequencing of the full-length LC sequence. The sensitivity of the RACE-RepSeq approach is achieved thanks to the inherent high Ig transcripts in PC, accounting for up to 70% of total mRNA.^31^ Consequently, even a very small fraction of monoclonal PCs, otherwise undetectable by conventional methods (including cytometry or DNA sequencing) is theoretically detectable using RACE-RepSeq.^32^ Accordingly, an IGLV clone was detected in 82% of BM analyzed, including 13 patients in which conventional methods did not allow the detection of an IGLV clone or detected an unrelated IGVK clone. This represents a 16% increase in the clone detection rate, compared to that reported in a study of 87 patients from the Mayo Clinic.^11^ Obviously, the detection of an IGLV clone in the BM does not mean it is the causative LC. But the combination of restricted usage of IGLV genes, the unique IGLJ domain (IGLJ3*02) found in 100% of POEMS syndrome LC sequences and the specific pattern of mutations revealed by full-length LC sequencing may increase the diagnostic sensitivity. When we added the LC sequences of POEMS syndrome of the present work with the series of Abe *et al*.^6^ and Martin *et al*.,^5^ the H40→N substitution in the FR2 region of IGLV1-40 sequences was found in 9/10 (90%) patients and the T38→P/A substitution in the CDR1 of IGLV1-44 sequences was found in 21/27 (78%) sequences. This appears highly specific to POEMS syndrome sequences since such mutations were almost never observed in sequences from AL amyloidosis or other PC disorders (~ 0.03% in a total of 124 IGLV1-40 and IGLV1-44 sequences) and none but one had a full PVNWYQ stretch found in IGLV1-44 sequences of POEMS syndrome but not associated with the recurrent IGLJ3*02 domain. It would have been interesting to analyze retrospectively the clinical symptoms of this unique AL amyloidosis patient with a clonal IGLV1-44 containing the PVNWYQ stretch (sequence DQ165740 in supplemental Table 1) since POEMS syndrome may be mistaken for an AL amyloidosis with neuropathy.^1^ Collectively it seems that these recurrent mutations are highly specific to POEMS syndrome and may be a reliable diagnostic marker. As an example, patient 20 presented with a polyneuropathy, elevated VEGF, 158 mg/l of λ free LC in the serum and a plasmacytoma but a κ^+^ monoclonal plasmocytosis in the BM and no λ LC clone found by Sanger sequencing in the BM. We considered that the BM plasmocytosis was not related to the POEMS syndrome and he was treated with irradiation. The treatment was unsuccessful and he subsequently received a high dose chemotherapy with autologous stem cell transplantation. Retrospectively, thanks to RACE-RepSeq approach, we found in the BM an IGLV1-40 monoclonal sequence with the H40→N substitution associated with the IGLJ3*02 domain. The dissemination of this λ PC clone probably explains the ineffectiveness of irradiation. Conversely, patient 27 presented with an isolated plasmacytoma, a discrete elevation of λ-LC PCs in the BM and a circulating monoclonal IgAλ. This patient was treated with irradiation and is still in clinical response after 8 years. We retrospectively performed RACE-RepSeq on the BM and detected an IGLV clone but derived from the IGLV3-1 germline gene, which was not reported in POEMS syndrome, confirming the lack of association between the BM clone and the POEMS syndrome. We have also confirmed in this study that some patients with POEMS syndrome have more than one monoclonal Ig and that full-length LC sequencing may help unveil the involved clone in such situation (p9). Finally, this combination of restricted IGLV usage and specific pattern of mutations can shed light on the mysterious pathophysiology of POEMS syndrome. The origin of the high level of VEGF, involved in many clinical features of the disease, is still controversial but appears to be directly correlated to the level of the monoclonal LC.^3^ Some studies have suggested that VEGF could be produced by monoclonal PCs in plasmacytoma^33^ or by both monoclonal and polyclonal PCs in BM.^34^ However, this hypothesis was refuted in a recent study showing that BM PCs of POEMS patients do not produce more VEGF-A as compared to MGUS ou MM PCs.^30^ Given the low level of BM infiltration by monoclonal PCs in POEMS patients, these controversies will likely be resolved by next generation cytometry analysis or single cell transcriptomic. One hypothesis is that the monoclonal λ LC could act as a paracrine or endocrine factor, that could activate the secretion of VEGF-A in cells/tissues. The restricted structure of monoclonal LCs from POEMS syndrome argues for a “LC-Target” interaction. Interestingly, when the PVNWYQ stretch was submitted to Blastp analysis,^35^ we found a 100% homology with the HSP90α chaperone protein known to be implicated in the regulation of angiogenesis through HIF1 chaperoning and to be a ligand for CD91 in its secreted form.^36,37^ Thus, we hypothesize that the λ monoclonal LC in POEMS syndrome could drive the secretion of VEGF by a mechanism of molecular mimicry with HSP90α.

In conclusion, using a new highly sensitive method of RNA repertoire sequencing (RACE-RepSeq), we confirmed the restricted usage of IGLV and IGLJ genes in POEMS syndrome, found a putative new germline gene (IGLV1-36) and highlighted peculiar patterns of mutations that could help to confirm the diagnosis and will likely shed new lights in the pathophysiology of this complex disease. Although it deserves to be confirmed with largest cohorts and with adequate controls, the sensitivity of RACE-RepSeq seems to outcompete other methods to detect small PC clones. The application and the benefits of RACE-RepSeq in other MGCS or in the detection of minimal residual disease in PC disorders in general deserve to be investigated.

## Acknowledgments

This work is support by Ligue Nationale contre le Cancer du Limousin and Fondation Française pour la Recherche contre le Myélome et les Gammapathies monoclonales. SB, DL and FA are supported by the French Ministry of Research ‘Plan maladies rares’. MVA is supported by Fondation ARC pour la Recherche sur le Cancer. We would like to acknowledge the use of the Boston University ALBase, supported by HL68705, in this work.

## Authorship Contributions

SB performed, analyzed experiments and wrote the manuscript. AS and MA collected and analyzed data and reviewed the manuscript. DL and FA collected clinical data. MVA drafted the manuscript. VJ, MC, MF, VP, FB critically reviewed the manuscript. CS and AJ supervised this study, analyzed experiments and wrote the manuscript.

## Patient protections

This study has received DC 2008-111 approval for retention and treatment of biological samples and CNIL (DR 211392) and CCTIRS (N°1158) approval for additional data processing.

