## Supplemental figure 1 for "Full-length immunoglobulin high-throughput sequencing reveals specific novel mutational patterns in POEMS syndrome"

A

| Patient Id | IGLV gene | Clone % |
|---|---|---|
| p2 | IGLV1-44 | 100% |
| p4 | IGLV1-44 | 38.63% |
| p5 | IGLV1-44 | 91.42% |
| p7 | IGLV1-44 | 91.2%* |
| p8 | IGLV1-44 | 91.3%* |
| p9 | IGLV1-44 and IGLV3-1 | 2.607% and 27,78% |
| p10 | IGLV1-44 | 43.62% |
| p11 | IGLV1-44 | 0.482% |
| p12 | IGLV1-44 | 4.591% |
| p13 | IGLV1-44 | 1.545% |
| p14 | IGLV1-44 | 41.95% |
| p15 | IGLV1-44 | 63.2% |
| p16 | IGLV1-44 | 49.77% |
| p17 | IGLV1-44 | 53.19% |
| p18 | IGLV1-44 | 84.89%* |
| p19 | IGLV1-40 | 54.98% |
| p20 | IGLV1-40 | 8.2% |
| p22 | IGLV1-40 | 59.32% |
| p23 | IGLV1-40 | 6.67% |
| p24 | IGLV1-40 | 71.93% |
| p25 | IGLV1-36 | 59.16% |
| p26 | IGLV1-36 and IGLV2-14 | 12.24% and 21.72% |
| p27 | IGLV3-1 | 89.3% |
| p28 | IGLV3-1 | 82.32% |
| p29 | IGLV3-1 | 2.569% |
| p30 | IGLV3-25 | 4.42% |

B

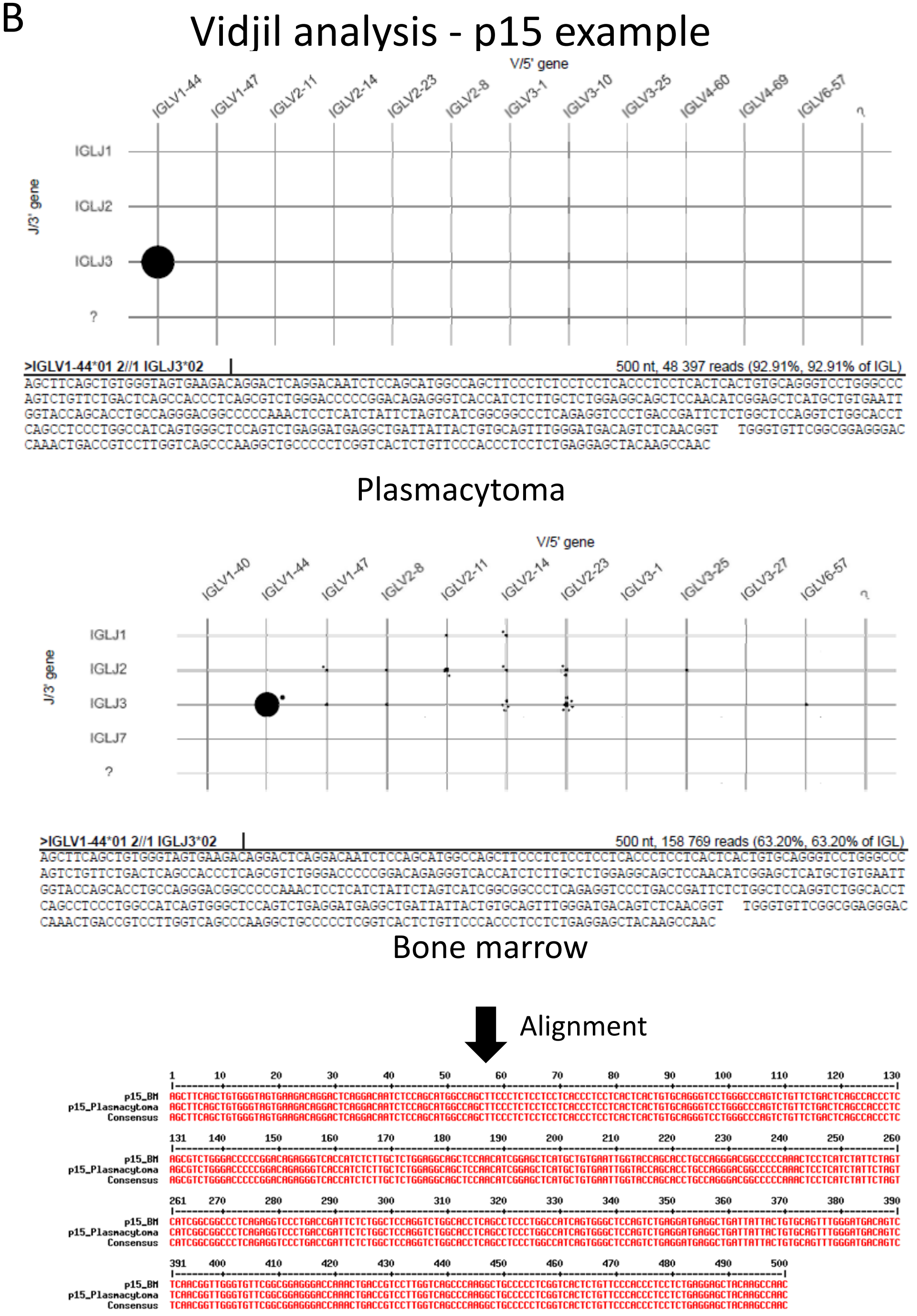


Supplemental Figure 1 : Analysis of the λ LC repertoire of patients with POEMS syndrome using Vidjil tool. (A) Percentage of the clonal λ LC sequences in the λ repertoire obtained with RACE-RepSeq and using vidjil tool for analysis (www.vidjil.org). All sequences were obtained from bone marrow samples except in * corresponding to plasmacytoma biopsies. (B) Example of vidjil tool analysis for patient 15 (p15). Clonal sequences from the bone marrow and the plasmacytoma are identical using multalin tool (http://multalin.toulouse.inra.fr/multalin/).
