## Supplemental figure 2 for "Full-length immunoglobulin high-throughput sequencing reveals specific novel mutational patterns in POEMS syndrome"

| | FR1 | CDR1 | FR2 | CDR2 | FR3 | CDR3 |
|---|---|---|---|---|---|---|
| EU599333 | QSVLTQPPS-ASGTPGQRVTISCSGG | SSNI----GSNT | VTWYQQLPGTAPKLLVY | TN-------N | QRPSGVP-DRFSGSK--SGTSASLAISGLQSEDEADYHC | AAWDDSLNG |
| DQ165743 | SPVLTQPPS-ASETPGQRVTISCSGS | SSNI----GSNT | VNWYQHLPGTAPKFLIY | AN-------N | QRPSGVP-DRFSGSK--SGTSASLAISGLQSEDEADYFC | ASWDDSLNG |
| EF589496 | QSVLTQPPS-ASGTPGQRVTISCSGS | ISNI----GSNT | VNWYQQLPGTAPKLLMY | SN-------D | QRPSGVP-ERFSGSK--SGTSVSLAISGLQSEDEADYYC | AAWDDSLNG |
| AF490941 | QSVLTQPPS-ASGTPGQRVTISCSGS | SSNI----GSNT | VNWYQQLPGTAPKFLIY | AN-------N | ERPSGVP-DRFSGSK--SDTSAALAISGLQSDDEADYYC | AAWDDSVNG |
| A42193 | QSVLTQPPS-ASGTPGQRVTISCSGS | SSNI----GSNV | VTWYQQLPGTAPKLLIY | TN-------N | QRPSGVP-GRFSGSK--SGTSASLAVSGLQSEDEADYYC | ATWDDS-VN |
| AF128428 | ---------------GQRVTISCSGS | SSNI----GSNT | VNWYQKLPGTAPKLLIY | ST-------- | QRPSGVP-GRFSGSK--SGTSASLAVSGLQSEDEADYYC | ATWDDS-VN |
| AF054638 | QSVLTQPPS-ASGTPGQRVTISCSGS | NSNI----GINI | VNWYQQVPGRAPKLLIY | SN------NN | QWPSGVP-DRFSGSK--SGTSASLAISELQSEDEGDYYC | ASWDDTMD- |
| AF128430 | --------------PGQRVTISCSGN | DSNI----AINS | VNWYQHLPGSAPKLLIY | SH-------- | QRPSGVA-DRFSGSK--SGTSASLAISGLQSEDEADYYC | AAWDDSLHG |
| EF589458 | QSVLTQPPS-ASGTPGQSITISCSGS | SSNI----GSNS | VTWYQHLPGTAPKLLIY | SV------NN | HRPSGVP-DRFSGSK--SGTSXSLAISGLQSEDEANYYC | AAWDDSLNG |
| AF054640 | QSVLTQPPS-ASGTPGQRVTISCSGS | RTNI----ASNT | VNWYQHLPGMAPKLLIY | TN-------D | QRPSGVP-ARFSGSK--SGTSASLAIIGLQSEDEADYYC | GAWDGSLSG |
| DQ165740 | SYELTQPPS-TSGAPGQRVTISCSGS | SSSI----GSN**P** | **VNWYQ**HLPGTAPKFLIY | AN-------- | QRPSGVP-DRISGSK--SGTSASLGISGLQSEDEADYYC | AAWDDRLNG |
| DQ165738 | SYELTQPPS-ASETPGQRVTISCSGS | SSNI----GSNT | VNWYQHLPGTAPKFLIY | AN-------N | QRPSGVP-DRFSGSK--SGTSASLAISGLQSEDEADYFC | ASWDDSLNG |
| EF589532 | QSVLTQPPS-ASGTPGQRVTISCSGR | SSNI----GSNT | VHWYQHVPGAAPKLLIY | SN------DN | QRPSGVP-DRFSGSK--SGTSASLAISGLQSEDEADYFC | ASWDDSLNG |
| AF115348 | QSVLTQPPS-ASGTPGQRVIISCSGS | SSNI----GRNT | VNWYLHLPGTAPKLVIY | SN-------D | ERPSGVP-DRFSGSK--SGTSASLAISRLQSEDEAHYYC | ATWDDSPNA |
| DQ165750 | QSVLTQPPS-ASETPGQRVTISCSGS | SSNI----GSNT | VNWYQHLPGTAPKFLIY | AN-------- | QRPSGVP-DRFSGSK--SVTSASLAISGLQSGDEAHSYC | AAWDDSLRG |
| DQ165748 | QSVLTQPPS-ASETPGQRVTISCSGS | SSNI----GSNT | VNWYQHLPGTAPKFLIY | AN-------N | QRPSGVP-DRFSGSK--SGTSASLAISGLQSEDEADYFC | ASWDDSLNG |
| AY701712 | ------------GTPGQRVTISCSGS | SSNI----GSNT | VNWYQQLPGTAPKLLIY | NN-------N | QRPSGVP-DRFSGSK--SGTSASLAISGLQSEDEADYFC | ASWDDSLNG |
| 05-086 | QSVLTQPPS-ASGTPGQRVTISCSGS | SSNI----GSNN | VNWYQQLPGMAPKLLIY | NI------NN | QRPSGVP-DRFSGSK--SGTSASLAINGLQSEDEADYYC | ASWDDSLN- |
| 06-098 | QSVLAQPPS-ASASPGQRVTVSCSGA | ISNI----GSNS | VYWYQQLPGTAPKLLIY | SD-------R | QRPSGVP-DRFSGSK--SGTSASLAIGGLQSEDEADYHC | VAWDDSLNA |
| AF462674 | QSVLTQPPS-ASGAPGQTVTISCSGR | TSNI----GSF**A** | VAWYQQVPKAGPKLLIF | RD-------S | HRLPGVP-DRFSGSK--SGTSASLAISGLQSEDEAHYHC | AAWDDSLNG |
| EF589490 | QSVLTQPPS-ASGTPGQRVTISCSGS | SSNI----GSNT | VNWYQHLPGTAPKFLIF | SN-------N | QRPSGVP-DRFSGSK--FGSSASLAISGLRSEDEADYYC | SSWDAGLNG |
| AF124167 | QSELTQPPS-ASGTPGQRVTISCSGS | SSNI----GSIT | VTWYQHLPGAAPKLLIY | SN-------D | QRPSGVP-DRFSGSK--SGTSASLAISGLQSEDEADYYC | AAWDDSLNG |
| 06-085 | QSVLAQPPS-ASASPGQRVTVSCSGA | ISNI----GSNS | VYWYQQLPGTAPKLLIY | SD-------R | QRPSGVP-DRFSGSK--SGTSASLAISGLQSVDEAHYYC | SAWDDRLNG |
| AF462684 | QSVLTQPIS-ASGTPGQRVSISCSGG | YYNI----RTNT | VHWYQQLPGTAPKLLIY | NN-------D | HRLPGVP-DRFSGSK--SGTSASLAISGLQSEDEAHYHC | -AAWDDSLNG |
| DQ165735 | QSVLTQPPS-ASGTPGQWVTISCSGG | NSNI----GTN**P** | VTWYQLLPGTAPKLLIY | DS-------- | QRPSGVP-DRFSGSK--SGPSASLAIGGLLSEDEADYYC | -ATWDDSLNG |
| EF589442 | QSVVTQPPS-ASGTPGQRVTISCSGS | SSNI----GSNT | VNWYQRLPGTAPKVLIY | SN------DH | QRPSGAP-DRFSGSK--SGTSASLVIRGLQSEDEADYSC | AAWDDSLSG |
| DQ165745 | QSVLTQPPS-ASETPGQRVTISCSGS | SSNI----GSNT | VNWYQHLPGTAPKFLIY | AN-------- | QRPSGVP-DRFSGSK--SDTSASLAISGLQAEDEADYFC | AAWDESLNG |
| EF589398 | QSLLTQPPS-ASGTPGQRVTISCSGR | RSNI----GSNT | VTWYQQLPGTAPKLLIY | TN------DN | QRPSGVP-DRFSGSK--SGTSASLAISGLQSEDEADYFC | ASWDDSLNG |
| 01-031 | QSVLTQPPS-ASGTPGQRVTISCSGS | SSNI----GSNT | VNWYQHLPGTAPKLLIY | SN-------N | LRPSGVP-DRFSGSK--SGTSGSLAISGLQSADEGDYYC | AAWDDSLNG |
| 02-005 | QSVLTQPPS-ASGTPGQRVTISCSGS | SSNI----GRNT | VNWYQQLPGTAPKFLIY | SN-------D | ERPSGVP-DRFSGSK--SGTSASLAISGLQSEDEADFYC | AAWDDSLTG |
| 06-082 | QSVLTQPPS-ASGTPGQGVTISCSGR | SSNI----GRNT | VNWYQKLPGTAPKFLIY | SN-------N | QRLSGVP-DRFSGSK--SGTSASLAIGGLQSADEATYYC | -ASWDDSLNG |
| EF589547 | QSVLTQPPS-ASGTPGQRVFVSCSGS | SSNI----GSNT | VNWYQHLPGTAPKLLIY | IT-------D | QRPSGVP-DRFSGSK--SGTSASLAISGLQSEDEADYYC | AAWDDSLDG |
| EU599332 | QSVLTQPPS-ASGTPGQSVTISCSGS | SSNI----GRNT | VNWYQHLPGTAPKLLIY | SN-------N | QRPSGVP-DRFSGSK--SGTSASLAITGLQSEDEADYYC | AVWDDSLNG |
| DQ165742 | SYELTQPPS-ASETPGQRVTISCSGS | SSNI----GSNT | VNWYQHLPGTAPKFLIY | AN-------- | ERPSGVP-DRFSGSK--SGTSASLAISGLRSEDEADYFC | AAWDDSLNG |
| EF589557 | QSVLTQPPS-ASGTPGQRVAISCSGS | NSNI----GRNS | INWYQHVPGTPPKLLIF | TN------NN | QRPSGVP-DRFSGSK--SGTSASLAISGLQSEDEADYFC | ASWDDSLNG |
| EF589503 | QSVLTQAPS-ASGTPGQTVTISCSGS | RSNI----GSNT | VNWYQQVPGTAPTFLIY | SN-------S | QRPSGVP-DRFSGSK--SGSSASLAIRGLQSQDEAFYYC | SSWDDNLNG |
| AY701714 | ------------GTPGQRVTISCSGG | SSNI----GINT | VDWYQHLPGTAPKLLIY | NN-------- | QRPSGVP-DRFSGSK--SGTSASLAISGLQSDDEADYYC | AAWDDSLNG |
| 05-109 | QSVLTQPPS-ASGTPGQRVTISCSGS | ISNI----RSNT | VNWYQQLPGTAPKVLIY | SN------NN | QRPSGVP-DRFSGSK--SGTSASLAIIGLQSEEEADFYW | ASWDDSLD- |
| DQ165737 | QSVLTQPPS-AYETPGQRVTISCSGS | SSNI----GSNT | VNWYQHLPGTAPKFLIY | AN-------- | QRPSGVP-DRFSGSK--SETSASLAISGLQSEDEADYFC | AAWDDSLNG |
| AF115347 | QSVLTQPPS-ASGTPGQRVTISCSGS | NSNI----GSNS | VNWYQHLPGTAPKLLIY | SQ------SN | QRPSGVP-DRFSGSK--SGTSASLAISGLQSEDEADYFC | ASWDDSLNG |
| EF589554 | QSVLTQPPS-ASGAPGQTITISCSGS | SSNI----GINT | VCWYQQYPGTAPKLLIY | TN-------D | QRPSGVP-DRFSGSR--SGTSASLAISGLQSEDEAEYYC | AAWDDSLNG |
| DQ165747 | QSVLTQPPS-ASETPGQRVTISCSGS | SSNI----GSNT | VNWYQHLPGTAPKFLIY | AN-------- | HRPSGVP-DRFSGSK--SGTSASLAISGLQSEDEADYYC | AAWDDILDG |
| AY701711 | ------------GTPGQRVTISCSGS | SSNI----GSNT | VNCYQQLPGTAPKLLIY | TN-------N | QRPSGVP-DRFSGSK--SGTSASLAISGLQSEDEADYFC | ASWDDSLNG |
| AF124170 | QSVHNQPPS-ASGTPGQRVTISCSGS | RTNI----ASNT | VNWYQHLPGIAPKLLIY | TN------DN | QRPSGVP-DRFSGSK--SGT-ASLAINGLQSEDEAYYYC | ATWDDIMNG |
| AY701709 | ------------GTPGQRVTISCSGS | RSNI----DANT | VNWCQHLPGTAPKLLIL | SN-------- | QRPSGVP-DRISGSK--SGTSASLAISGLQSEDEAHYYC | AAWDDRLNG |
| DQ240234 | QSVLTQPSS-ASGTPGQRATISCSGT | YSNV----GFNS | VNWYQLLPGTAPKLLIY | AT------SN | QRPSGVP-DRFSGSK--SGTSASLAISGLQSEDEADYYC | ATWDDSVNG |
| AF124166 | QSVLTQPPS-ASGTPGQRVTISCSGS | NSNI----GINI | VNWYQQVPGRAPKLLIY | SN-------N | ERPSGVP-DRFSGSK--SGTSASLAISGLQPEDEADYYC | VAWDDLLKA |
| DQ165734 | QSVLTQPPS-ASETPGQRVTISCSGS | SSNI----GSNT | VNWYQHLPGTAPKFLIY | AN-------- | QRPSGVA-DRFSGSK--SGTSASLAISGLQSEDEADYYC | AAWDDSLHG |
| EF589510 | QSVLTQPPS-ASGAPGQRVTISCSGS | SSNL----GSNN | VNWFQQLPGAAPKLLIY | TN------NN | QRPSGVP-DRFSGSK--SGTSASLAISGLQSEDEADYFC | ASWDDSLNG |
| EU599334 | QSVLTQPPS-ASGTPGQRVTLFCSGS | SSNI----GSNT | VQWYQHLPGTAPKLLIY | TN-------D | QRPSGVP-DRFSGSK--SGTSASLAITGLQSEDGADYYC | ATWDDSLNG |
| DQ165744 | SPVLTQPPS-ASETPGQRVTISCSGS | SSNI----GSNT | VNWYQHLPGTAPKFLIY | AN-------- | QRPSGVP-DRFSGSK--SGTSASLAIGGLQSEDEADYFC | SAWDDSLNG |
| EU599319 | QSVLTQPPS-ASGTPGQRVTISCSGS | SSNI----GGNV | VNWYQQLPGMAPKVLIH | SQ------NN | QRPSGVP-DRFSGSK--SGTSASLAISGLQSEDEADYFC | ASWDDSLNG |
| EF589497 | QSMLTQPPA-ASGTPGQRVSISCSGS | SSNI----GSNT | VNWYQQLPGTGPKLLIF | SS-------N | QRAPGVP-DRFSGSK--SDTSASLAISGLQSEDEADYYC | ASWDDSVQG |
| EF589474 | QSVLTQPPS-ASGTPGQRVTISCSGS | TSNI----GTNT | VNWYQHVPGAAPQLLIY | TN-------D | QRPSGVP-DRFSGSK--SGTSASLAISGLQSEDEADYYC | AAWDDSLNG |
| AF490942 | ------------GTPGQRVTISCSGS | RSNI----GENT | VSWYQQLPGTAPKFIIF | SN-------D | QRPSGVP-DRISGSK--SGTSASLAISGLQSEDEADYHC | AAWDDSLHG |
| AF054639 | QSVLTQPPS-ASGTPGQRVTISCSGS | SSNI----GSNT | VTWYQHLPGAAPKLLIY | SN-------D | QRPSGSL-TDSLAPS--LAP- |  |
| AF128431 | ---------------GQRVTISCSGS | XSNI----AINS | VNWYQHLPGSAPKLLIY | SH-------- | QRPSGVP-DRFSGSK--SGTSASLAISGLQSVDEAHYYC | SAWDDRLNG |
| AF054641 | QSVLTQPPS-ASGTPGQRVSISCSGS | TSNI----GSNT | VNWYLHLPGTAPKLLIY | SN------NN | HRPSGVP-DRFSGSK--SGTSASLAISGLQSEDEANYYC | AAWDDSL-- |
| EF589484 | QSVVTQPPS-ASGTTRQRVTISCSGG | SSNV----GSNT | VNWYQHLPGTAPKLLIY | SG-------N | QRPSGVP-DRFSGSK--SGTSASLAISGLQSEDEADYYC | AAWDDSLDG |
| EU599331 | QSVLIQPPS-ASGTPGQRVTISCSGS | SSNI----GINN | VNWYQHLPGTAPKLLIY | TN-------N | QRPSGVP-DRFSASK--SGTSXSLAISGLQSDDEGDYYC | AVWDDSLNG |
| AF320833 | QSVLTQAPS-ASGTPGQRVTISCSGS | SSNI----GSNT | ANWYQHLPGTAPKLLIY | TN-------S | ERPSGVP-DRFSGSK--SGTSASLAISGLQSEDEADYYC | SSWDESLNG |
| EF589384 | QSVLTQPPS-ASGTPGQRVTISCSGS | SSNL----GSNT | VNWYQHLPGSAPKLLIY | SN-------N | QRPSGVP-DRFSGSK--SGTSASLAISGIQSEDEAVYYC | AAWDDSLDG |
| DQ165741 | SYELTQPPS-ASETPGQRVTISCSGS | SSNI----GSNT | VNWYQHLPGTAPKFLIY | AN-------- | QRPSGVP-DRFSASK--SGTSASLAISGLQSEDESNYYC | AAWDARLKG |
| DQ165739 | SYELTQPPS-ASETTGQRVTISCSGS | SSNI----GSNT | VNWYQHLPGTAPKFLIY | AN-------N | QRPSGVP-DRFSGSK--SGTSASLAISGLQSEDEADYFC | ASWDDSLNG |
| EF589533 | QSVLTQPPS-TSGTPGQRVIISCSGT | RSNI----GSNI | VNWHQQFPGTAPKLLIY | SD------DN | QRPSGVP-DRFSGSK--SGTSASLAISGLQSEDEADYFC | ASWDDSLNG |
| AF320843 | QSVLTQPPS-ASGTPGQRVTISCSGS | SSNI----GRNT | VHWYQQLSGTAPKFLIF | NT-------Y | QRPSGVP-DRFSGSK--SGTSASLAISGLQSEDEADYYC | AAWDDSLNG |
| EF589556 | QSVLTQPPS-ASGTPGQRVTISCSGS | NSNI----GINI | VTWYQQVPGRALKLLIY | SN-------N | DRPSGVA-DRFAGSK--SDTSASLAISGLQSDDEADYYC | ATWDDNLNE |
| DQ165749 | QSVLTQPPS-ASETPGQRVTISCSGS | SSNI----GSNT | VNWYQHLPGTAPKFLIY | AN-------- | QRPSGVA-DRFSGSK--SGTSASLAISGLQSEDEADYYC | AAWDDSLHG |
| AY701713 | ------------GTPGQRVTISCSGS | SSNI----GGNT | VSWYQQLPGTAPKLLIN | SN-------N | QRPSGVP-DRFSGSK--SGTSASLAISGLQSEDEADYFC | ASWDDSLNG |
| EF589479 | QSVLTQPPS-ASGTPGQRVTISCSGS | NFNV----GRNT | VQWYQQVPGSAPKLLIY | AD------DV | QRPSGVP-ERFSGSK--SDTSASLAISGLQSEDEADYYC | AAWDDSVNG |
| EF589520 | QSVLTQPPS-ASGNPGQRVTISCSGT | ISNI----GSNT | VNWYQHLPGTGPKLLIF | SN-------N | QRLSGVP-DRFSGSK--SGTSASLAISGLHSEDEAAYSC | ATWDDRLSG |
| 05-108 | QSELTQPPS-ASGTPGQRVTISCSGS | SSNV----GSNV | VNWYQQLPGTAPKILIY | SN-------T | QRPSGAP-DRFSGSR--SGTSASLAISGLRSEDEADYFC | AAWDDTLNG |
| EF589456 | QSVLTQPPS-ASGTPGQVVTMSCSGS | SSNI----GGNT | VAWYQHLPGAAPRLLVY | SN-------S | QRPSGVP-DRFSGSK--SGTSASLAISGLQSEDEADYYC | AAWDDSLNG |
| 0610195A | ESVLTQPPS-ASGVPGQSVIISCSGS | SSNI----GRNT | VNWYQQVPGAAPKLLVY | SN-------N | QWPSGVP-DRFSGSK--SGTSASLAISGLHSEDEADYFC | ATWDDSLDG |
| AF124168 | QSVQTHPTS-ASGTPGQRVIISCSGS | SSNI----GRNT | VNWYLHLPGTAPKLVMY | SN-------D | QRPSGVP-DRFSGSK--SVTSASLAISGLQSEDEAHYYC | AAWDDSLRG |
| AF462685 | QSALTQPPS-ASGTPGQRVTISCYGS | SSNI----GRTT | VNWYQQVPGTGPKFLIY | NN-------N | QRPSGVS-DRFSGSK--SGTSASLAISGLQSDDEADYYC | AAWDDSLNG |
| DQ165736 | QSVLTQPPS-ASETPGQRVTISCSGS | SSNI----GSNT | VNWYQHLPGTAPKFLIY | AN-------N | LRPSGVP-DRFSASK--SGASASLAISGLRSEDEADYYC | ATWDDSLS- |
| EF589489 | QSVLTQPPS-ASGTPGQRVTISCSGS | RSNI----GGDS | VNWYKHLPGSGPKLLIF | GT-------D | VRTSGVP-ERFSGSK--SGTSASLAISGLQSEDEGDYFC | AAWDDNLNG |
| DQ165746 | QSVLTQPPS-ASETPGQRVTISCSGS | SSNI----GSNT | VNWYQHLPGTAPKFLIY | AN-------N | QRPSGVP-DRFSGSK--SGTSASLAISGLQSEDEADYFC | ASWDDSLNG |
| AY701710 | ------------GTPGQRVTISCSGS | SSNI----GSNT | VNWYQQLPGTAPKLLIY | SN-------N | QRPSGVP-DRFSGSK--SGTSASLAISGLQSEDEADYYC | AAWDDSLNG |
| DQ098823 | ----------ASGTPGQRVTISCSGS | TSNI----GRNT | VNWYQQLPGTAPKLLIY | SN-------D | QRPSGVP-DRLSGSK--YATAASLAINGLQSEDEADYYC | AVWDDSLNG |
| DQ098788 | -SVLTQPPS-ASGTPGQRVTISYSGS | SSNI----GSIT | VNWHQQLPGPAPKLLIY | GN-------D | HRPSGVP-DRFSGSK--SGTSASLAISGLQSEDEGTYYC | AAWDDSLN- |
| DQ098789 | QSVLTQPPS-ASGAPGQRVSISCSGS | SSNI----GDNN | VNWYQQLPGTAPRLLMY | SN-------N | QRPSGVP-DRFSGSK--SGTSASLAISGLQSEDEADYYC | TSWDGRLKG |
| 1006259B | ESVLTQPPS-ASGTSGQRVTISCSGS | SSNL----GSNQ | VNWYRHLPGTAPKLVIY | SD-------S | QRPSGVP-DRISASK--SGTSASLAISGLQSEDESDYYC | ASWDDSLDG |
| 1006259E | ESVLTQPPS-ASGTPGQRVTISCSGS | GSNI----GSHT | VNWYHQFPGTAPKLLIY | RN-------D | QRPSGVP-DRFSGSK--SGTSASLAISGLQSEDEADYYC | AAWDDSLDG |
| AY271357 | QSVLTQPPS-ASGTPGQRVTISCSGS | SSNI----GINS | VNWYQQFPGKAPKLVVV | LN-------N | QRPSGVA-DRFSGSK--SGTSASLAISGLQSEDEADYYC | ATWGGSLNA |
| Per1 | QSVLTQPPS-ASGTPGQRVTISCSGS | RSNI----GSNT | VNWYHQLPGTAPRLLIY | AN-------N | QRPSGVP-DRFSGSK--SGTSASLAISGLQSEDEADYYC | AAWDDSLNG |
| Per2 | QSVLTQPPS-ASGTPGQRVTISCSGS | SSNI----GSNT | VNWYQQLPGTAPKFVIY | SN-------N | QRPSGVP-DRFSGSK--SGTSASLAISGLQSEDEADYHC | ASWDDSLNG |
| Per3 | QSVLTQPPS-ASGTPGQRVTMSCSGS | SSNI----GPNT | VNWYHQLPGTAPKLLIY | IN-------N | QRPSGVP-DRFSGSK--SGTSASLAISGLQSEDEADYFC | AAWDDSLNG |
| Per4 | QSVLTQSPS-ASGTPGQRVTISCSGS | RSNI----GTNT | VNWYHQLPGTAPKLLIY | NN-------N | QWPSGVP-DRFSGSK--SGTSASLAISGLQSEDEADYYC | AAWDDSLNG |
| Per5 | QSVLAQPPS-ASGTPGQRVTISCSGS | SSNI----GRNT | VNWYQQLPGTAPKLLIY | IN-------H | QRPSGVP-DRFSGSK--SGTSASLAISGLQSEDEGDYYC | AAWDDTLNG |
| Per6 | QSVLTQPPS-ASGTPGQRVTISCSGS | RNNI----GSNS | VTWYQQLPGTAPKLLIY | SN-------N | QRPSGVP-DRFSGSK--SGTSGSLAISGLQSDDEADYYC | AAWDDSLNG |

```
Per7           QSFLTQSPS-ASGTPGQRVTISCSGS SSNI----GSNT VNWNQQLPGTAPKLLIY ND-------L QRPSGVP-DRFSGSK--SGTSASLAISGLQSEDEADYYC ATWDDSVNG
Per8           QSVLTQPPS-TSGTPGQRVTISCSGS SSNI----ETNT VNWYQQLPGTAPKLVMH TN-------N QRPSGVP-DRFSGSR--SGTSASLAIGGLQSEDEADYFC AAWDDNLNG
Per9           QSVLTQPPS-ASGPPGQRVTISCSGS RSNI----GGNS VVWYQQLPGTAPNLLIF NT-------N ERPSGVP-DRFSGSK--SGTSASLAISGLQSEDEADYFC AAWDDTLTG
Per10          QSELTQPPS-VSGTPGQRITISCSGS SSNL----GHNS VHWYQHLPGTAPKLLIF SN-------S QRPSGVP-DRFSGSK--SGTSASLAISGLQSGDDVDYYC AAWDDSLNG
Per11          QSVLTQPIS-ASGTPGQRVSISCSGG YYNI----RTNT VHWYQQLPGTAPKLLIY NN-------D QRPSGVP-DRFSGSK--SGPSASLAIGGLQSEDEADYYC ATWDDSLNG
Per12          QSVLTQPPS-ASGAPGQTVTISCSGR TSNI----GSFA VAWYQQVPKAGPKLLIF RD-------S QRPSGVP-DRFSGSK--FGSSASLAISGLRSEDEADYYC SSWDAGLNG
Per13          QSVLTQPPS-ASGTPGQRVTISCSGS SSNI----GSNT VNWYQQLPGTAPKLLMY RN-------D QRPSGVP-DRFSGSK--SGSSASLAISGLQSEDEADYYC AAWDDSLNG
Per14          QSALTQPPS-ASGTPGQRVTISCYGS SSNI----RTNT VNWYQQVPGTGPKFLIY NN-------N QRPSGVS-DRFSGSK--SGTSASLAISGLQSDDEADYYC AAWDDSLNG
Com2           QSVLTQAPS-ASGTPGQRVTISCSGS SSNI----GSNT ANWYQHLPGTAPKLLIY TN-------S QRPSGVP-DRFSGSK--SGTSASLAISGIQSEDEAVYYC AAWDDSLDG
Com3           QSVLTQPPS-ASGTPGQRVTISCSGS NSNI----GSNS VNWYQHLPGTAPKLLIY SQ-------S QRPSGVP-DRFSGSR--SGTSASLAISGLQSEDEAEYYC AAWDDSLNG
Com4           QSVLTQPPS-ASGTPGQRVSISCSGS TSNI----GSNT VNWYLHLPGTAPKLLIY SN-------N QRPSGVP-DRFSGSK--SGTSASLAISGLQSEDEADYYC AAWDDSLDG
Com6           QSVLTQPPS-ASGTPGQRVTISCSGS SSNI----GRNT VHWYQQLSGTAPKFLIF NT-------Y DRPSGVA-DRFAGSK--SDTSASLAISGLQSDDEADYYC ATWDDNLNE
AL1            QSVLTQPPS-ASGTPGQRVIISCSGG TSNI----GDNS VVWYQQLPGTAPKLLIY SN-------D QRPSGVA-DRFSGSK--SGTSGSLAISALQSEDEADYYC SSWDDSLHG
AL2            QSVLTQPPS-ASGTPGQRVTISCSGS SSNI----ASNS VTWYQHLPGTAPKLLIY SN-------N QRPSGVP-DRFSGSK--SGTSASLAISGLQSEDEADYYC GVWDDSVTG
IGLV1-44       QSVLTQPPS-ASGTPGQRVTISCSGS SSNI----GSNT VNWYQQLPGTAPKLLIY SN-------N QRPSGVP-DRFSGSK--SGTSASLAISGLQSEDEADYYC AAWDDSLNG

                           FR1              CDR1           FR2             CDR2                   FR3                     CDR3
AF462673       QSVLTHPPS-VSGAPGQRVTISCTGS RSNIG---SGYE IHWYQQLPGEAPQLLIY GD-------T NRPSGVP-DRFSGSK--SGTSASLAITGLQAEDEADYHC QSYDISRSG
AF320834       QSVLTQPPS-VSGAPGQWVTISCTGS RSNIG---AGFD VHWYQQLPGSAPKLLIY AN-------I YRPSGVP-DRFSGSK--SGTSASLAITGLQPEDEANYYC QSYDNNPNT
EF589558       QSVLTQPPS-VSGAPGQRVTISCTGS SSNIG---AGYY VNWYQQFPGTAPRLLIY GN-------I NRPSGVT-DRFSGSK--SGTSASLAIAGLQAEDEADYYC HSYDNSLSA
EF589450       QSVLTQPPS-VSGAPGQRVTISCTGS SSNIG---AGYD VHWYQQLPGTAPKLLIY AN-------N NRPSGVP-DRFSASK--SGTSASLAITGLQAEDEADFYC QSYDSSLGG
EU599322       QSVLTQPPS-VSGAPGQRVTISCTGS RSNIG---AGYH VHWYQQLPGTAPKLLIY AD-------T NRPSGVP-DRFSGSK--SGTSASLAITGLQAEDEAEYYC QSYDTNLV-
EF589512       QSVLTQPPS-VSGAPGQRVTISCTGS TSNIG---ARYD VHWYQQLPGAAPKLLIY GN-------T NRPSGVP-DRFSGSK--SGTSASLAITGLQAEDEADYYC QSYDNSLSG
EF589542       QSLLTQPPS-VSGAPGQGVTISCTGS NSNIG---TGYD LHWYQQLPGTAPKLLIY GN-------T NRPSGVP-DRFSGSK--SGTSAYLAITGLQAEDEADYYC QTYDSSVSA
AF320844       QSVLTQPPS-VSGAPGQRVTISCTGS SSNIG---AGID VHWYQQVPGTAPHLLIY DN-------I NRPSGVP-DRFSGSK--SATSASLAITGLQADDEADYYC QSYDSSLSG
AF490938       ------XPX-VSGAPGQRVTISCTGS SSNLG---AGYD VHWYQQLPGTAPKIVIY GN-------N IRPSGVP-DRISGSK--SGTSASLAITGLQAEDEADYYC QSYDRSG--
EF589403       QSVLTQPPS-VSGAPGQRVTISCTGS SSNIG---AGYE VHWYQQPPGTAPKLLMY GN-------T NRPSGVP-DRFSGSK--SGSSASLAITGLQAEDESDYYC QSYDSSMSG
EF589431       QSVLTQPPS-VSGAPGQRVTISCTGS SSNIG---ARYD VHWYQHLPGTAPKLLIY AN-------N NRPSGVP-DRFSGSK--SGTSASLAITGLQAEDEALYYC QSYDSSLSD
P01703         QSVLTQPPS-VSGAPGQRVTISCTGS SSNIG---AGNH VKWYQQLPGTAPKLLIF HN-------N --------ARFSVSK--SGSSATLAITGLQAEDEADYYC ---------
DQ098786       QSVLTQPPS-VSGAPGQRVTISCTGS SSNIG---AGYD VHWYQQVPGTAPKLLIY GN-------K NRPSGVP-DRFSASK--SGTSASLAITGLQADDEADYYC QSYDSSLR-
DQ098812       ----TQPPS-VSGAPGQRVTISCTGS RSNIG---AGYD VHWYQQLPGTAPKLLIY GN-------T NRPSGVP-NRFSGAK--SGTSASLAIAGLQAEDEGDYYC QSYDNSLSG
DQ098814       QSVLTQPPS-VSGAPGQRVTISCTGN KSNIG---AGYD VHWYQLIPGTAPKRLIY GN-------V NRLSGVP-DRFSGSK--SGTSASLAITGLQAEDEADYYC QSYDTSVNG
AY180392       ----------------QRVTISCTGS SSNIV---EGHD VHWYQQFPGKAPKLLIY GN-------N NRPSGVP-DRFSGSK--SGTSASLAITGLQAEDEADYYC QSYDSSLSF
AL3            QSVLTQPPS-VSGAPGQRVTISCTGS SSNIG---AGYD VHWYQQIPGTAPKLLIY LN-------T NRPSGVP-DRFSGSK--SDTSASLAITGLQAEDEADYYC QSYDNSLSG
AL4            QSVLTQPPS-VSGAPDQRVTISCTGT SSNIG---AHYD VHWYQHLPGTAPKLLIY GN-------N NRPSGVP-DRFSGSK--SDISASLAITGLQAEDEGDYFC QSYDRSLSG
IGLV1-40       QSVLTQPPS-VSGAPGQRVTISCTGS SSNIG---AGYD VHWYQQLPGTAPKLLIY GN-------S NRPSGVP-DRFSGSK--SGTSASLAITGLQAEDEADYYC QSYDSSLSG
```

Supplemental Figure 2: Deduced amino-acid sequences of monoclonal IGLV1-44 and IGLV1-40 λ light chain variable domains of AL amyloidosis or other plasma cell disorders patients compared to germline sequences according to IMGT numbering. Per1 to Per14 from Perfetti et al[26], com2 to com6 from Comenzo et al[27], AL1 to AL4 personal data, all other sequences from the Boston university AL-Base database[28]. A and P mutations in position 38 for IGHVL1-44 and N in position 40 for IGHVL1-40 are in red underlined.
